# RNA Mango-based Sensors for Lead

**DOI:** 10.1101/2025.05.07.652743

**Authors:** Annyesha Biswas, Saurja DasGupta

## Abstract

Lead (Pb^2+^) toxicity poses a serious threat to human health and remains a global concern; therefore, there is a critical need for the development of easy-to-use and cost-effective tools for the rapid monitoring of Pb^2+^. In this study, we demonstrate the potential of the RNA Mango aptamer as a sensitive and selective sensor for Pb^2+^. Our findings reveal that trace amounts of Pb^2+^ induce the formation of the G-quadruplex motif in RNA Mango, which facilitates dye binding and activates fluorescence. A detailed investigation of the fluorescence properties of RNA Mango with three different dyes, TO1-Biotin, TO3-Biotin, and Thioflavin-T, in the presence of Pb^2+^ shows that RNA Mango has the highest binding affinity for Pb^2+^ in combination with TO1-Biotin, with a *K*_D_ value as low as ∼100 nM. In the presence of Pb^2+^, RNA Mango has sub-micromolar affinity for all three dyes, showing the tightest binding to TO1-Biotin (*K*_D_ ∼40 nM). Mango lead sensors detect low nanomolar concentrations of Pb^2+^ with limits of detection of 2 – 16 nM, which are significantly lower than its allowable limit in drinking water. RNA Mango exhibits remarkable selectivity toward Pb^2+^ and can detect lead in tap water samples. This work reports a new class of simple and inexpensive fluorescence-based sensors for lead and expands the repertoire of RNA-based lead sensors.

## Introduction

Heavy metal contamination is a persistent global threat to environmental and human health. Among these metals, lead (Pb^2+^) is of particular concern due to its high toxicity and harmful physiological effects even at low exposure levels. Industrial processes such as the mining and smelting of lead ores,^1^ production of leaded gasoline^2^, and the manufacturing of lead-acid batteries^3^ and lead-containing pigments^4^ release large amounts of lead into the environment. Importantly, chronic exposure to low amounts of lead, primarily through contaminated drinking water,^5^ industrial waste,^6^ and deteriorating infrastructure such as lead-based pipes,^7^ has been linked to cognitive deficits,^8^ developmental disorders,^9^ cardiovascular diseases,^10^ and kidney dysfunction.^11^ Pb^2+^ toxicity poses a heightened risk to children, where exposure is associated with neurodevelopmental impairments, behavioral disorders, and reduced attention spans.^12,13^ Given these health risks, regulatory agencies such as the U.S. Environmental Protection Agency (EPA) and the World Health Organization (WHO) have set stringent limits for lead in drinking water at 15 parts per billion (72 nM) and 10 parts per billion ∼46 nM), respectively.^14,15^ Ensuring compliance with these regulations requires sensitive and reliable detection methods.

Traditional analytical techniques for Pb^2+^ detection include atomic absorption spectrometry, inductively coupled plasma mass spectrometry, anodic stripping voltammetry, and X-ray fluorescence spectroscopy.^16^ These methods allow highly sensitive Pb^2+^ detection; however, the requirements for expensive and elaborate instrumentation, trained personnel, and complex sample preparation make them less suitable for rapid, on-site monitoring. Organic scaffolds, like small molecule chelators, macrocycles, or even metal-organic frameworks (MOFs), have been used for fluorescence-based and colorimetric detection of Pb^2+^.^17–19^ Although these chemical sensors are capable of detecting Pb^2+^ without sophisticated instrumentation, they are not sufficiently selective and often fail to detect trace amounts of Pb^2+^.

To address these limitations, there is increasing interest in developing biosensors that offer rapid, cost-effective, and portable lead detection with minimal technical expertise. Biosensors generally outperform chemical sensors in selectivity and sensitivity metrics, presumably due to more sophisticated molecular recognition by biomolecules.^20,21^ Nucleic acid-based biosensors detect Pb^2+^ by leveraging structural changes upon Pb^2+^ binding.^22,23^ For example, Pb^2+^ binding to RNA-cleaving DNAzymes induce the formation of their catalytic conformation, which results in the cleavage of an RNA strand.^24^ Pb^2+^-induced cleavage of a fluorescently-labeled RNA substrate spatially separates the fluorophore on the RNA from the quencher on the DNAzyme, generating a fluorescent ‘On’ signal. A similar strategy can be used to create colorimetric DNAzyme-based sensors by conjugating them to gold nanoparticles.^25^ G-quadruplex-based biosensors exploit the Pb^2+^-induced folding of guanine-rich sequences into a G-quadruplex. G-quadruplexes are typically formed in the presence of monovalent cations like Na^+^ and K^+^.^22^ Pb^2+^, due to its similar ionic radius to K^+^ and twice the charge, facilitates the formation of a stable G-quadruplex structure at lower concentrations. G-quadruplex-based sensors exploit the structural change from single-stranded to G-quadruplex in different ways.^26^ Pb^2+^-induced quadruplex formation may bring the two DNA termini modified with a fluorophore and quencher in close proximity, resulting in a fluorescence ‘Off’ signal.^27^ Certain G-quadruplex sequences exhibit peroxidase activity upon binding hemin and appropriate reactants like ABTS to generate colored, chemiluminescent, or fluorescent products.^28^ Therefore, the activity of these quadruplex-hemin DNAzymes has also been used as a Pb^2+^ sensing platform. DNA-based electrochemical sensors convert G-quadruplex formation to an electrochemical signal, which is detected by techniques such as cyclic voltammetry.^29^ However, the requirement for nucleic acid modification with fluorophores/quenchers or nanoparticles, pH-sensitive chemical reactions, or expensive electrochemical assemblies pose significant challenges to the practical application of these biosensors for Pb^2+^ detection.

A new strategy that leveraged Pb^2+^-induced G-quadruplex formation in the fluorogenic RNA aptamer, Spinach, generated the first RNA-based biosensor for Pb^2+^.^30^ Stabilization of the Spinach G-quadruplex by trace amounts of Pb^2+^ allows its cognate dye, DFHBI, to bind to this platform, activating strong and stable green fluorescence. Spinach demonstrated moderate binding affinity for Pb^2+^ (*K*_D_ ∼1.3 μM) but high sensitivity with a limit of detection (LOD) of 6 nM. This system consists of just the RNA aptamer and its cognate dye, where the fluorescence signal is a result of non-covalent dye binding. Therefore, fluorogenic RNA aptamers offer a simple and efficient platform for Pb^2+^ detection, addressing key limitations related to covalent modifications of nucleic acids with dyes or nanoparticles, optimal reaction conditions for colorimetric or chemiluminescent detection, and elaborate electrochemical assemblies. Despite the creation of numerous G-quadruplex-containing fluorogenic RNA aptamers since its development, the Spinach sensor remained the only RNA-based biosensor for Pb^2+^. In this study, we expand the repertoire of RNA-based Pb^2+^ sensors by evaluating alternative fluorogenic RNA aptamers for enhanced binding affinity to Pb^2+^, increased sensitivity, and improved selectivity. We identify the Mango RNA aptamer as an excellent candidate for creating Pb^2+^ sensors and describe its Pb^2+^-sensing properties in combination with three dyes: TO1-Biotin and TO3-Biotin, the cognate dyes for this aptamer, and Thioflavin-T, a sequence agnostic, but G-quadruplex-specific dye.^31–34^ A detailed characterization of the fluorescent properties of the Mango aptamer with these dyes in the presence of Pb^2+^ allowed us to develop the most sensitive RNA-based Pb^2+^ sensors to date.

## Results and Discussion

### RNA Mango exhibits strong fluorescence in the presence of Pb^2+^

Since the development of the Spinach sensor, other G-quadruplex-containing fluorogenic RNA aptamers such as Broccoli, Mango, Peach, and Beetroot have been artificially evolved (Fig. S1). ^35–38^ The Broccoli (Fig. S1A) and Beetroot (Fig. S1B) aptamers were isolated from random RNA libraries by virtue of their ability to bind to DFHBI-1T and DFHO and exhibit green and yellow fluorescence, respectively (Table S1).^35^ The Mango aptamer (Fig. S1C) was similarly obtained for binding to Thiazole Orange derivatives, TO1-Biotin, and TO3-Biotin^36^ and Peach, an aptamer related to Mango, was selected for binding to TO3-Biotin.^38^. The G-quadruplex motif in each aptamer, stabilized by 100 mM K^+^ in the folding buffer, serves as the dye binding platform and is consequently key to the aptamer’s fluorescent properties. To assess whether these G-quadruplex-containing RNA aptamers can be used to detect Pb^2+^, we examined the fluorescence properties of Broccoli, Mango, Peach, and Beetroot in the presence of their respective dyes, both in the absence and presence of 1 µM Pb^2+^ (Figs. 1 and S2).

**Fig. 1.**
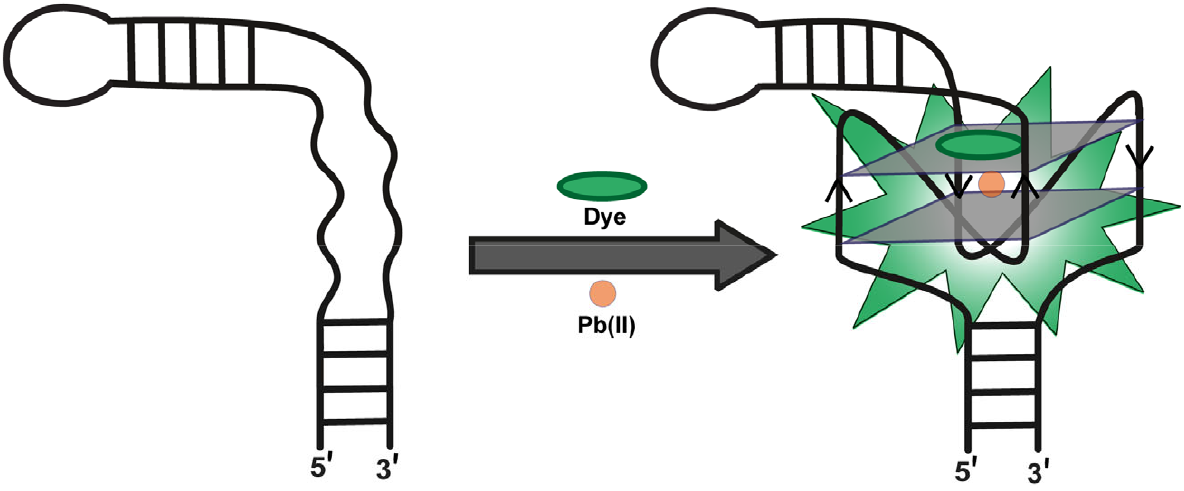
Schematic illustration of a fluorogenic RNA-based lead sensor. The unstructured region (shown by curved lines) in the RNA aptamer folds into a G-quadruplex structure in the presence of Pb^2+^ and binds to the fluorophore, triggering a fluorescence signal.

Although all four aptamers use G-quadruplexes to bind their cognate dyes, only RNA Mango in combination with TO1-Biotin exhibited a significant fluorescence enhancement (∼8-fold) in the presence of 1 μM Pb^2+^ (Fig. S2A). The Peach/TO3-Biotin combination also showed fluorescence enhancement upon the addition of Pb^2+^, however, this enhancement was less pronounced (3-fold) (Fig. S2B). No noticeable enhancement in the fluorescence was observed for the Broccoli/DFHBI-1T and Beetroot/DFHO aptamer-dye pairs (Figs. S2C and S2D). The fluorescence enhancement exhibited by Mango/TO1-Biotin in the presence of 1 µM Pb^2+^ was 33-fold, 308-fold, and 1449-fold higher compared to Peach/TO3-Biotin, Broccoli/DFHBI-1T, and Beetroot/DFHO, respectively (Fig. 2A). These results suggested that among the aptamers tested, Mango was the most promising candidate for a Pb^2+^sensor.

**Fig. 2.**
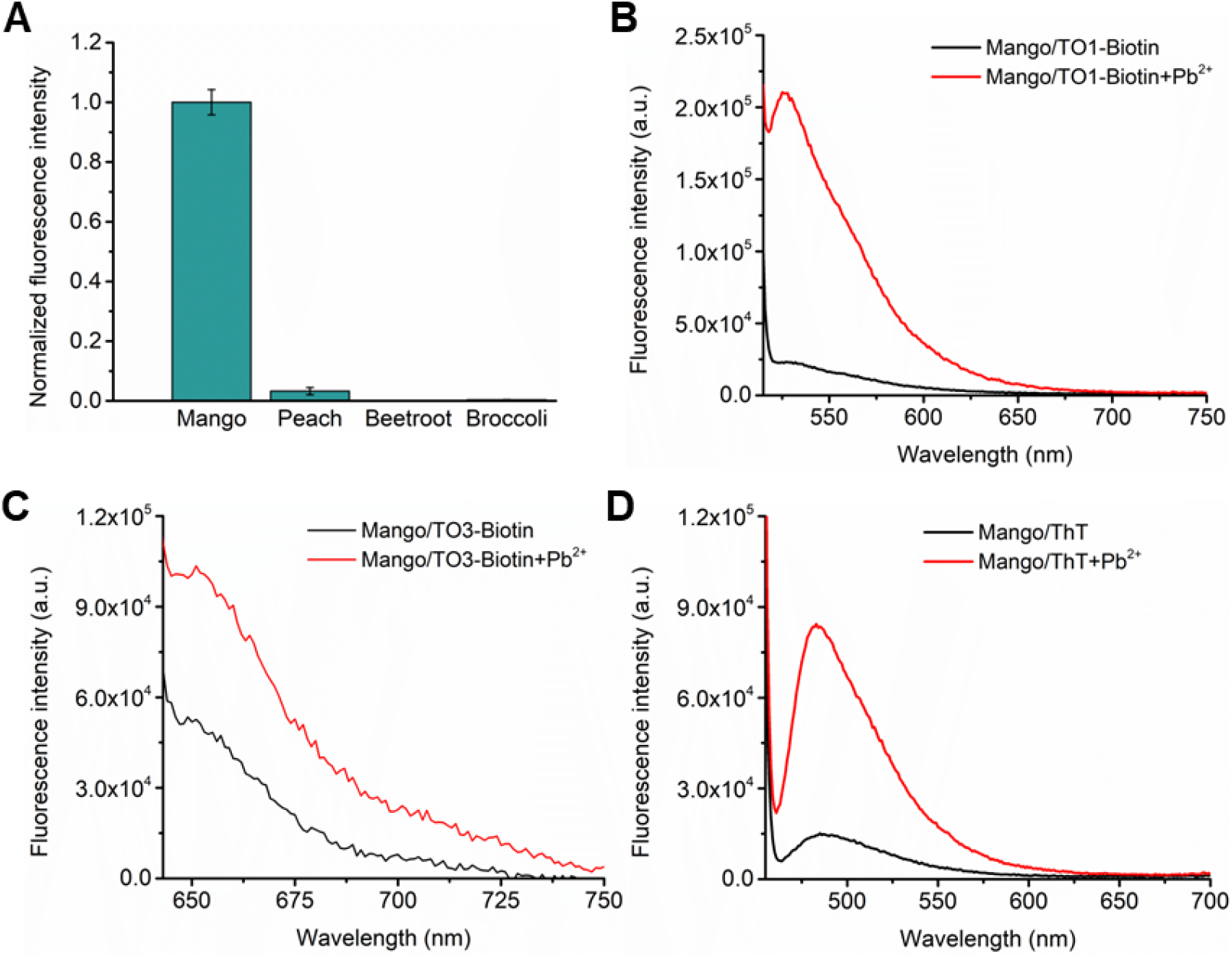
RNA Mango exhibits fluorescence enhancement in the presence of Pb^2+^. (A) Fluorescence enhancement in different RNA aptamers/dye combinations upon the addition of 1 μM Pb^2+^. (B)-(D) Fluorescence spectra of (B) Mango-TO1-Biotin, (C) Mango-TO3-Biotin, and (D) Mango-ThT, in the absence (black) and presence of 1 μM Pb^2+^ (red). Mango-TO1-Biotin and Mango-ThT showed significant fluorescence enhancement with Pb^2+^. Concentrations of the RNA and dye were 300 nM and 3 μM, respectively. Experiments were performed at pH 8 and 5 mM Mg^2+^. Error bars in (A) correspond to the standard deviation of three independent experiments.

As Mango showed the highest increase in fluorescence in the presence of Pb^2+^, we studied its fluorescence properties with TO3-Biotin and Thioflavin-T (ThT), dyes that have been previously reported to bind Mango in addition to TO1-Biotin.^39,40^ While TO3-Biotin is a cognate dye of Mango and Peach aptamers, ThT exhibits green fluorescence upon specifically binding to G-quadruplex structures in a context-independent manner. (Table S1).^31–34^ Similar to TO1-Biotin, Mango showed fluorescence enhancements of ∼2.1-fold and ∼5-fold in the presence of TO3-Biotin and ThT, respectively upon Pb^2+^ addition (Figs. 2C, D). This ability of Mango to exhibit fluorescence enhancements with all three dyes in the presence of just 1 μM Pb^2+^ highlights its robustness and versatility as a potential Pb^2+^ sensor.

### Pb^2+^-induced fluorescence activation of Mango is dependent on G-quadruplex stabilization

Stabilization of the G-quadruplex motif in Mango is central to its fluorescence activation by Pb^2+^ as this motif serves as the binding platform for the dye. RNA G-quadruplexes usually exist in the parallel topology and are characterized by a prominent positive peak at ∼260 nm and a prominent negative peak at ∼240 nm.^41^ We observed an enhancement of the positive peak at 263 nm upon the addition of 1 μM Pb^2+^ to the Mango aptamer in the absence of any dye, which further increased in intensity in the presence of 10 μM Pb^2+^ (Fig. 3A). This Pb^2+^-induced increase in CD intensity at ∼263 nm was accompanied by a decrease in intensity at ∼240 nm. These results provided direct evidence of the Pb^2+^-induced stabilization of the G-quadruplex motif in Mango that is independent of dye binding.

**Fig. 3.**
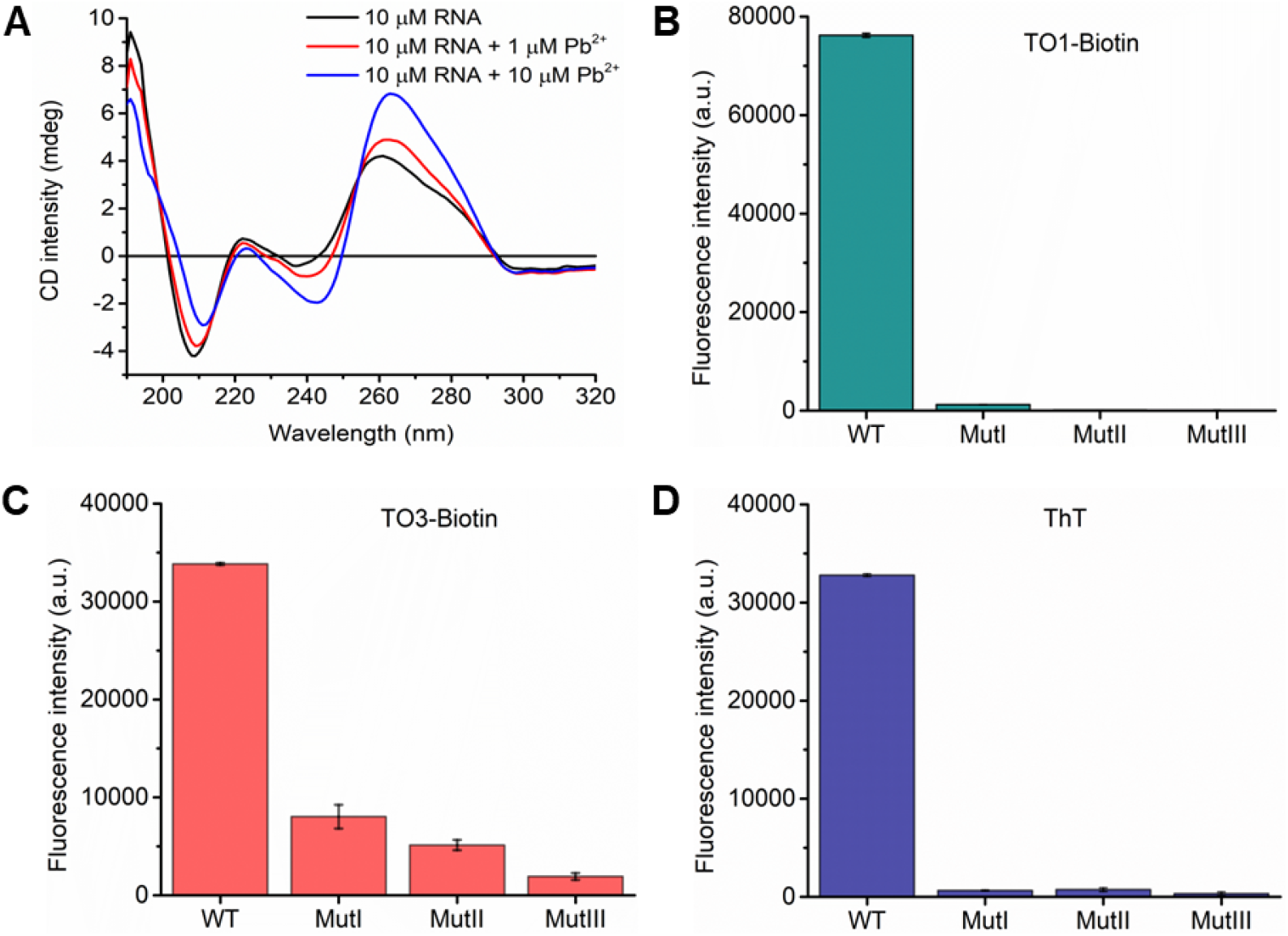
The G-quadruplex motif in RNA Mango is central to its ability to detect Pb^2+^. (A) CD spectra of the RNA Mango aptamer show an increase in the intensity of the positive peak at ∼260 nm and a decrease in the intensity of the negative peak at ∼240 nm after addition of 1 μM (red) and 10 μM (blue) Pb^2+^. This suggests that Pb^2+^ stabilizes the RNA Mango G-quadruplex. The spectrum in black was obtained without Pb^2+^. Experiments were performed at pH 8, 5 mM Mg^2+^, and 10 μM RNA. (B)-(D) Mutations to G-quadruplex nucleotides abolish Pb^2+^-induced Mango fluorescence with TO1-Biotin and ThT and significantly reduce fluorescence with TO3-Biotin. WT Mango shows strong fluorescence in the presence of Pb^2+^, but quadruplex mutants of the sensor, MutI, MutII, and MutIII, do not fluoresce over background with TO1-Biotin and ThT dyes. Detectable fluorescence exhibited by the quadruplex mutants with TO3-Biotin may be attributed to the intrinsic fluorescence of the dye in buffer (see Fig. S4B). Concentrations of RNA and dye were 100 nM and 1 μM, respectively. Error bars in (B)-(D) correspond to the standard deviation of three independent experiments.

To further investigate the effect of Pb^2+^-induced G-quadruplex formation on the fluorescence properties of Mango, we mutated the guanine residues critical for quadruplex formation to adenines (Table S2). Mutating these key guanine residues in the G-quadruplex resulted in a near-complete loss of fluorescence for TO1-Biotin and ThT (Fig. 3B, D). Although significantly reduced, some fluorescence was detected even in the quadruplex mutants of Mango when bound to TO3-Biotin (Fig. 3C), likely due to the higher intrinsic fluorescence of TO3-Biotin in buffer. Collectively, these results confirm that Pb^2+^-induced fluorescence activation of Mango is dependent on G-quadruplex formation.

### RNA Mango is a sensitive Pb^2+^ sensor

An ideal Pb^2+^ sensor should exhibit high sensitivity with a limit of detection in the low nanomolar range. To evaluate its sensitivity, we investigated Mango fluorescence with TO1-Biotin, TO3-Biotin, or ThT in response to increasing concentrations of Pb^2+^. The binding affinity of Mango to Pb^2+^ in the presence of each dye was determined by fitting the data to the Hill equation (Figs. 4A-C). The apparent *K*_D_ values for Pb^2+^ binding with Mango in the presence of TO1-Biotin, TO3-Biotin, and ThT dyes were calculated as 112.6 ± 30.7 nM, 294.8 ± 52.2 nM, and 157.0 ± 32.0 nM, respectively (Fig. 4D). These results revealed that sensors created with Mango/TO1-Biotin, Mango/TO3-Biotin, and Mango/ThT bind Pb^2+^ with 11.6-fold, 4.4-fold, and 8.2-fold greater affinities than the Spinach sensor (*K*_D_ ∼1.3 μM).^30^ Limit of detection (LOD) of the Mango lead sensor with TO1-Biotin, TO3-Biotin, and Thioflavin-T were calculated as 2.05 nM, 15.79 nM, and 3.89 nM (Figs. 4E, S3), respectively (based on 3σ/slope method, where σ is the standard deviation of the blank). This is consistent with their trend in their binding affinities for Pb^2+^. Sensors made with Mango/TO1-Biotin and Mango/ThT are the most sensitive RNA-based Pb^2+^ sensors reported to date. Importantly, the LOD values for all three Mango-based lead sensors are significantly lower than the acceptable limits for Pb^2+^ in drinking water.

**Fig. 4.**
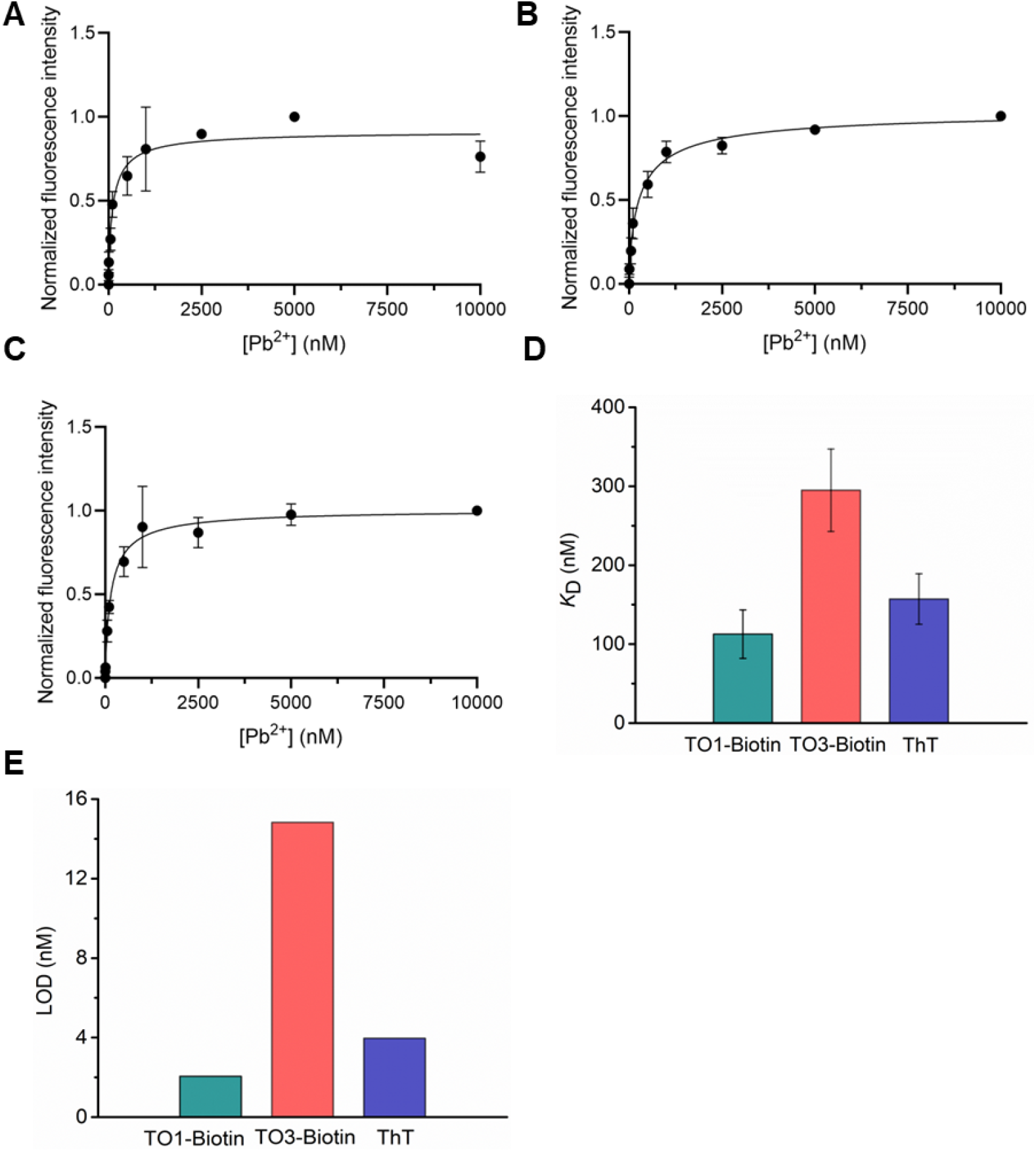
RNA Mango exhibits high sensitivity for Pb^2+^ detection. (A-C) Fluorescence signal exhibited by RNA Mango in response to increasing concentrations of Pb^2+^ with (A) TO1-Biotin, (B) TO3-Biotin, and (C) ThT. (D) RNA Mango binds to Pb^2+^ with high affinity in the presence of all three dyes, exhibiting *K*_D_ values of 100-300 nM. (E) RNA Mango is highly sensitive to low nanomolar concentrations of Pb^2+^ in the presence of all three dyes, with limits of detection (LOD) of 2-16 nM. Experiments were performed at pH 8 and 5 mM Mg^2+^. Concentrations of RNA and dye were 100 nM and 1 μM, respectively, with Pb^2+^ concentration in the range of 5 nM – 10 μM. Error bars correspond to the standard deviation of three independent experiments.

The increase in fluorescence signal in response to increasing concentrations of Pb^2+^ was directly imaged for all three Mango-based sensors (Fig. S4). While the Mango/TO1-Biotin and Mango/ThT sensors exhibited a clear, concentration-dependent enhancement of green fluorescence upon Pb^2+^ addition with negligible signal without Pb^2+^, the red fluorescence enhancement in the Mango/TO3-Biotin sensor was less discernible due to the high intrinsic fluorescence of the TO3-Biotin dye. Although Pb^2+^-induced fluorescence enhancement in the Mango/TO3-Biotin sensor can be clearly measured by fluorimetry, its inability to provide a distinct visual readout of this enhancement prevents its use as a point-of-care lead sensor in its current form. In contrast, the ease of Pb^2+^ detection by the Mango/TO1-Biotin and Mango/ThT combinations makes them attractive candidates for cheap, easy-to-use, point-of-care sensors for lead.

In addition to the sensors’ high affinity to Pb^2+^, the strong fluorescence signal in the presence of trace amount of Pb^2+^ could be due to tighter binding of the dye. To assess the affinity of RNA Mango to the three dyes in the presence of 1 μM Pb^2+^, we performed fluorescence titrations in the presence of increasing concentrations of TO1-Biotin, TO3-Biotin, and ThT (Fig. S5A-C). The apparent *K*_D_ values for dye binding were determined as 39.21 ± 7.64 nM for TO1-Biotin (Fig. S5A, D), 791.70 ± 72.20 nM for TO3-Biotin (Fig. S5B, D), and 192.60 ± 27.48 nM for ThT (Fig. S5C, D). The highest affinity of Mango to TO1 is consistent with the highest sensitivity and Pb^2+^ affinity exhibited by the Mango/TO1-Biotin sensor. In comparison. the apparent *K*_D_ value for DFHBI binding for the Spinach sensor in the presence of 10 μM Pb^2+^was determined as 1.14 ± 0.09 μM.^30^ Therefore, Mango-based lead sensors bind their dyes with at least 1.4-28.5-fold greater affinity than the Spinach sensor. Consequently, the Mango-based lead sensors require lower dye concentrations for Pb^2+^ detection.

### RNA Mango is a selective Pb^2+^ sensor

To investigate the selectivity of Mango sensors for Pb^2+^ detection, we measured their fluorescence in the presence of thirteen other metal ions that may be encountered in environmentally relevant samples (Fig. 5). Pb^2+^-specific fluorescence signal could be detected visually for the Mango/TO1-Biotin and Mango/ThT sensors; however, this was not possible for the Mango/TO3-Biotin sensor, which showed high background signal due to the intrinsic fluorescence of the TO3-Biotin dye (Fig. S6). In fluorimetric experiments, we observed strong fluorescence enhancement only upon Pb^2+^ addition with all three dyes; however weak signals were detected in the presence of Ca^2+^ and K^+^ with TO1-Biotin (Figs. 5A, B) and TO3-Biotin (Figs. 5C, D). In contrast, the Mango/ThT sensor showed weak but detectable fluorescence only when Ca^2+^ was present (Figs. 5E, F). These results are consistent with reports of G-quadruplex stabilization by Ca^2+^ and K^+^, with the lower signals reflective of much weaker binding than Pb^2+^.^30^ The Mango/TO1-Biotin sensor exhibited ∼20-fold and ∼25-fold higher signal for Pb^2+^ over K^+^ and Ca^2+^, respectively (Fig. 5B), whereas the Mango/TO3-Biotin sensor showed a ∼10-fold higher signal for Pb^2+^ relative to both Ca^2+^ and K^+^ (Fig. 5D). In contrast, the Mango/ThT sensor exhibited a 53-fold and 142-fold higher signal for Pb^2+^ compared to Ca^2+^ and K^+^, respectively (Fig. 5F). The superior selectivity for Pb^2+^, combined with the lowest limit of detection, makes Mango/ThT the best Mango-based lead sensor. The combined attributes of high binding affinity, excellent sensitivity, and strong selectivity for Pb^2+^ detection underscore the potential of Mango-based sensors as robust tools for the quantitative detection of Pb^2+^ at low concentrations.

**Fig. 5.**
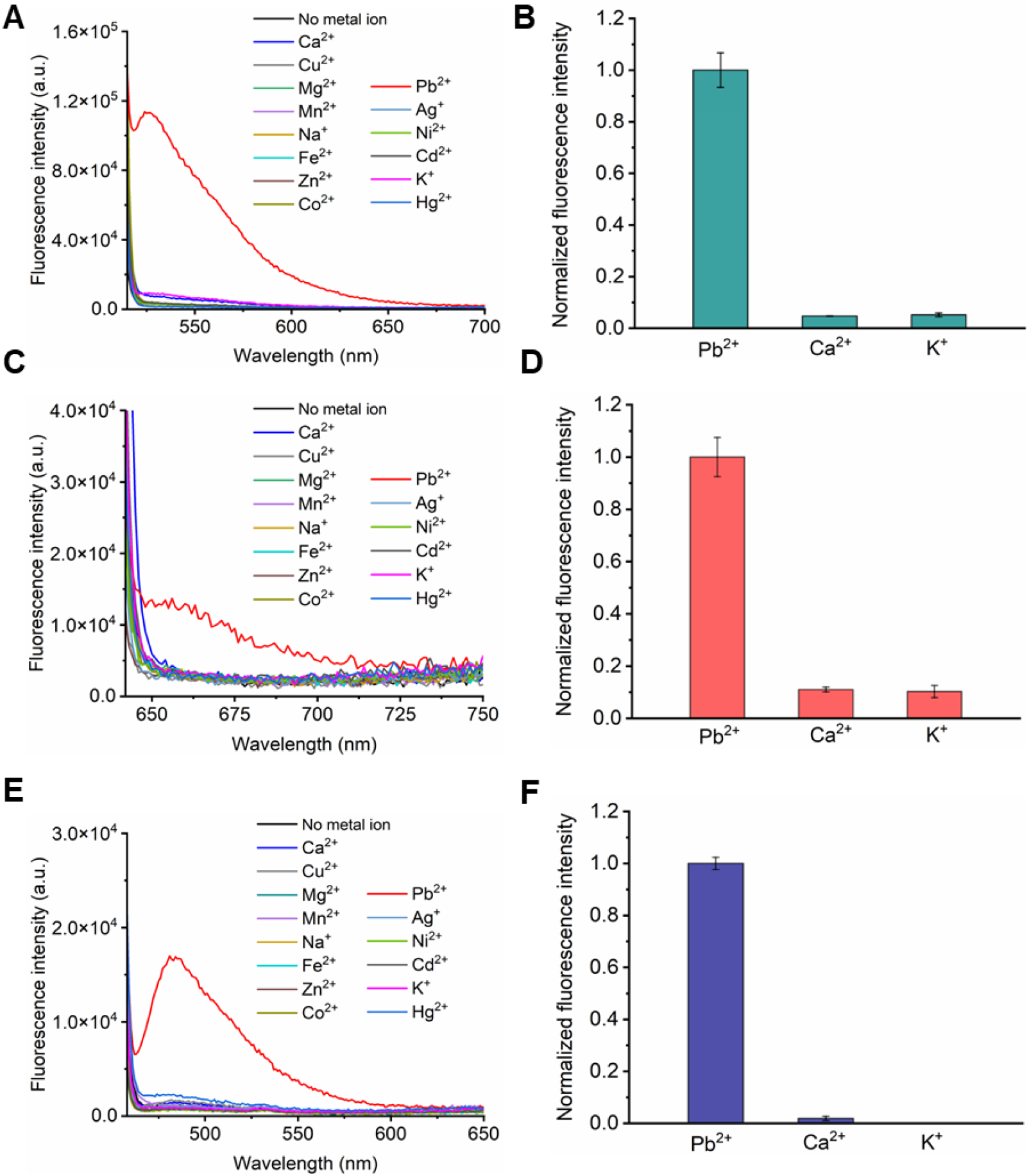
RNA Mango is selective for Pb^2+^. Fluorescence spectra of the Mango sensor with (A) TO1-Biotin, (C) TO3-Biotin, and (E) ThT in the presence of various metal ions. Relative fluorescence intensities of the Mango sensor in the presence of Pb^2+^, Ca^2+^, and K^+^ with (B) TO1-Biotin, (D) TO3-Biotin, and (F) ThT. Experiments were performed with 100 nM RNA, 250 nM dye, and 1 μM of metal ions at pH 8 and 5 mM Mg^2+^. Error bars in (B), (D), and (F) correspond to the standard deviation of three independent experiments.

### Mango detects Pb^2+^ in tap water

A reliable sensor must demonstrate functionality in practical, real-world applications. To evaluate this potential, we tested the applicability of the Mango lead sensors for detecting Pb^2+^ in tap water. Mango sensors emit fluorescence within minutes of encountering Pb^2+^, and previous studies have demonstrated that RNA-based sensors remain stable in tap water for days.^30^ Based on this, we assessed the performance of the Mango/TO1-Biotin and Mango/ThT sensors by spiking tap water samples with different concentrations of Pb^2+^. We observed an increase in fluorescence signal with Pb^2+^ concentrations ranging from 5 nM to 5 μM for TO1-Biotin (Fig. 6A) and ThT (Fig. 6B), indicating that the Mango sensor can detect Pb^2+^ in tap water. However, both Mango sensors showed modest background fluorescence in tap water samples, even in the absence of Pb^2+^. High concentrations of Ca^2+^ in the hard water samples may interfere with Pb^2+^ detection by Mango sensors; however, this can be remedied by several approaches as discussed in the next section.

**Fig. 6.**
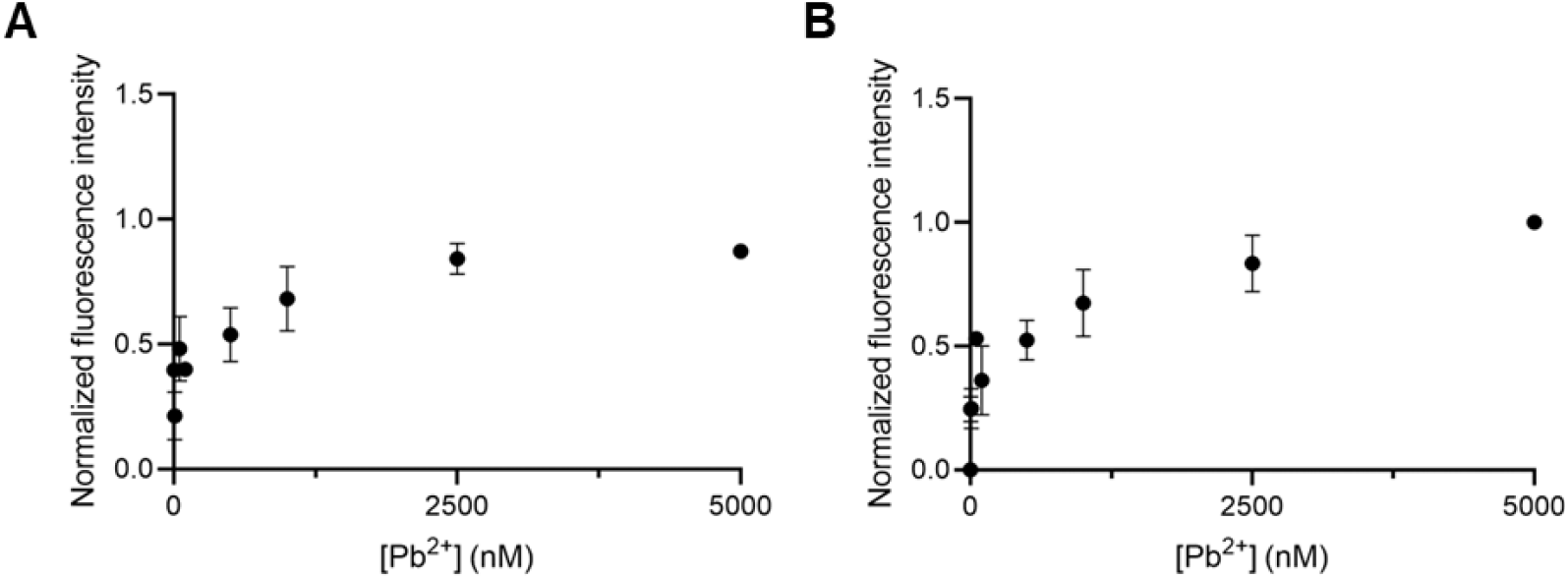
Mango lead sensors are functional in tap water. Fluorescence signal exhibited by RNA Mango in response to increasing concentrations of Pb^2+^ in tap water for (A) TO1-Biotin, and (B) ThT. Experiments were performed at pH 8 and 5 mM Mg^2+^. Concentrations of RNA and dye were 100 nM and 1 μM, respectively, with Pb^2+^ concentration in the range of 5 nM-5 μM. Error bars correspond to the standard deviation of three independent experiments.

### Conclusions

In this work, we created a new class of fluorescent biosensors for detecting Pb^2+^ based on the RNA Mango aptamer, thus expanding the analytical applications of RNA in heavy metal sensing. Notably, our sensors demonstrate excellent sensitivity with the best sensors exhibiting limits of detection below 1 part per billion (<1 ppb). The Mango sensors are able to discriminate between the Pb^2+^ and other metal ions, showing only a weak signal with K^+^ and Ca^2+^. The Mango/ThT sensor emerged as the most promising candidate exhibiting high affinity to its analyte, Pb^2+^ and its dye, ThT in the presence of Pb^2+^, and the highest selectivity among the Mango sensors. The versatility of Mango-based lead sensors is highlighted by their compatibility with dyes spanning different spectral regions, including TO1-Biotin and ThT in the green and TO3-Biotin in the red regions of the spectra, which allows users to select the dye best suited to their needs. Additionally, to detect lead, one simply needs to add the sample to the RNA/dye solution, eliminating the need for complex chemical reactions or sophisticated sensor assembly. Fluorescence signal can be observed in the presence of Pb^2+^ within minutes by simply shining light from a handheld UV lamp. These features make Mango lead sensors attractive for on-site detection of Pb^2+^.

However, to enable broad applicability in point-of-care devices, we are currently optimizing the concentrations of the Mango RNA aptamer and its fluorogenic dyes to reduce responsiveness to Ca^2+^, which, if present in real-world samples, may trigger weak false positives. Alternatively, water samples may be pre-treated with NH_4_OH to precipitate Ca^2+^ as Ca (OH)_2_, which can then be removed via filtration. A subsequent pH adjustment to ∼pH 8 with HCl will allow Mango sensors to function as intended. As Ca^2+^ is primarily present as Ca(HCO_3_)_2_ in hard water, boiling water samples will remove dissolved Ca^2+^ as CaCO_3_, allowing Pb^2+^ detection by Mango sensors. Another consideration for on-site detection is the potential to incorporate Mango sensors into inexpensive and portable point-of-care analytical devices. Toward this goal, we are developing paper-based analytical devices (PADs) and microfluidic chips for Pb^2+^detection that use the Mango sensors reported in this work.

## Experimental

### Materials

Tris-HCl (pH 8) buffer was purchased from Invitrogen. Metal salts Pb(NO_3_)_2_.3H_2_O, AgNO_3_, CdCl_2_.H_2_O, CuSO_4_.5H_2_O, FeSO_4_.7H_2_O, NiNO_3_.6H_2_O, ZnCl_2_, HgNO_3_.H_2_O, and Mn (CH_3_COO)_2_.4H_2_O were purchased from Sigma-Aldrich (ACS grade). MgCl_2_, NaCl, KCl, and CaCl_2_ were purchased as filtered solutions from Invitrogen. Dyes DFHBI, DFHBI-1T, and DFHO were purchased from Lucerna Technologies, TO1-Biotin and TO3-Biotin were purchased from Applied Biological Materials Inc. (abm), and Thioflavin-T (ThT) was purchased from Millipore Sigma. The details of all the dyes used in this work are listed in Supplementary Table S1. The RNA oligonucleotides used in this work are listed in Supplementary Table S2 and were purchased from Integrated DNA Technologies (IDT).

## Methods

### Instrumentation for fluorescence assays

Fluorescence emission was measured using a Fluorolog-3 spectrofluorometer equipped with a thermo-controller (Horiba Inc.) with a slit width of 2 nm for all the measurements. Excitation wavelengths of 510 nm, 637 nm, and 450 nm and emission ranges of 515-700 nm, 642-750 nm, and 455-600 nm were used for the fluorescence measurements for TO1-Biotin, TO3-Biotin, and Thioflavin T, respectively (See Supplementary Table S1). All measurements were done in triplicate at 25 °C. Data were plotted in Origin 2025 and GraphPad Prism 8.4.3.

### Identification of optimal aptamer-dye pairs for Pb^2+^ detection

To identify the optimal RNA-dye pairs for Pb^2+^ detection, we screened four fluorogenic RNA aptamers that use G-quadruplex motifs as their dye-binding platform. In these experiments, 300 nM RNA was heated in the presence 10 mM Tris-HCl (pH 8) at 90 °C for two minutes, followed by incubation with 5 mM MgCl_2_ (for the Mango) or 1 mM MgCl_2_ (for the Peach, Broccoli, and Beetroot aptamers) at 50 °C for 15 minutes. 3 μM dye (TO1-Biotin for Mango, DFHBI-1T for Broccoli, TO3-Biotin for Peach, or DFHO for Beetroot) was added to this solution and incubated at 37 °C for 10 minutes. Finally, 1 μM concentration of Pb(NO_3_)_2_ was added and incubated at 37 °C for 10 minutes. Fluorescence intensities for samples containing RNA and dye (‘blank’) were subtracted from the fluorescence intensities for samples containing RNA, dye, and Pb^2+^.

### Sensitivity and selectivity assays

Samples for sensitivity assays contained 1 μM dye, 100 nM RNA, and 5 nM -10 μM Pb^2+^. Binding affinities of RNA Mango for Pb^2+^ in the presence of the three dyes were determined by plotting normalized fluorescence (normalization was performed with respect to the highest signal) for each concentration of Pb^2+^ and fitting the data to the Hill equation: y = B_max_*x^h^ / (*K*_D_^h^ + x^h^), where B_max_ = highest fluorescence signal, *K*_D_ = dissociation constant, h = hill slope. The limit of detection (LOD) was measured from sensitivity assays using the 3σ/slope method (where σ represents the standard deviation of the 10 blanks). The fluorescence signal corresponding to the linear region of the binding plot was fitted into a simple linear regression model to obtain the slope. Samples for selectivity assays contained 100 nM RNA, 250 nM dye, and 1 μM metal ions (Pb^2+^ or non-Pb^2+^).

Sensitivity and selectivity assays were performed with Mango and each of the three dyes (TO1-Biotin, TO3-Biotin, or ThT). In addition to measuring fluorescence intensities using a fluorimeter, we imaged samples containing different Pb^2+^ concentrations (0.01, 0.1, 0.5, 1, 5, and 10 μM) or different metal ions at 1 μM by an Azure 600 imaging system using filters closest to the excitation and emission wavelengths of each dye. Samples containing TO1-Biotin, TO3-Biotin, and ThT were imaged under AzureRed (λ_ex_= 524 nm/λ_em_= 572 nm), AzureSpectra 650 (λ_ex_= 628 nm/λ_em_= 684 nm), or AzureSpectra 490 (λ_ex_= 472 nm/λ_em_= 513 nm), respectively.

### Dye-binding assays

5 nM -10 μM of each of the three dyes – TO1-Biotin, TO3-Biotin, or ThT were added to 100 nM RNA in the presence of 1 μM Pb^2+^ and incubated at 37 °C for 10 minutes. Fluorescence intensities at each dye concentration in buffer containing Pb^2+^, but in the absence of RNA, were subtracted from the fluorescence intensities observed for samples containing both dye and RNA. Binding affinities of the Mango lead sensor for each dye in the presence of Pb^2+^ were determined by plotting normalized fluorescence at each dye concentration and fitting the data to a Hill equation.

### Mutational studies on the RNA Mango G-quadruplex

We determined fluorescence intensities for three RNA Mango constructs (MutI, MutII, and MutIII; see Table S2) with mutations to their G-quadruplex regions. Fluorescence was measured for each mutant construct in the presence of TO1-Biotin, TO3-Biotin, or ThT. For each construct, the fluorescence signal without Pb^2+^ was subtracted from the fluorescence observed with Pb^2+^.

#### Detection of Pb^2+^ in tap water

To demonstrate the Pb^2+^ detection ability of the Mango sensors in a real-world sample, we added tap water samples spiked with 5 nM - 5 μM Pb^2+^ to the Mango sensors (100 nM RNA + 1 μM TO1-Biotin or ThT incubated at 37 °C for 10 minutes). The samples were further incubated at 37 °C for 10 minutes, and fluorescence intensities were measured as above. The average fluorescence intensity of 10 samples containing 100 nM RNA, 1 μM TO1-Biotin or ThT, and tap water was subtracted from each of the above Pb^2+^ spiked samples to calculate the fluorescence intensity solely due to Pb^2+^ in the spiked tap water samples.

#### Circular Dichroism (CD) spectroscopy

CD samples were prepared same way as in the case of fluorescence experiments, except that they contained 10 μM RNA Mango and did not contain any dye. CD measurements were carried out in a Jasco J-1500 CD spectrometer in the wavelength range of 180 to 320 nm, using a path length of 1 mm. Each CD spectra is an average of three scans measured at a rate of 50 nm min^-1^ and 1 nm interval. All measurements were taken at 25 °C. Data were smoothened and plotted in Origin 2025.

## Author contributions

A.B., and S.D. designed research; A.B. performed research; A.B., and S.D. analyzed the data; A.B., and S.D. wrote the paper.

## Data availability

Additional supporting data for this article have been included in the Electronic Supplementary Information file. This includes 1) Chemical structures of the dyes and the sequences of the oligonucleotides used in this work, 2) Secondary structures of the fluorogenic RNA aptamers tested and their fluorescence spectra with their corresponding dyes in the absence and presence of Pb^2+^, 3) Sensitivity plots of Mango sensor with TO1-Biotin, TO3-Biotin and ThT for Pb^2+^, 4) Images of the Mango lead sensors with increasing concentration of Pb^2+^ and with different metal ions, and 5) Binding curves for measuring the affinity of TO1-Biotin, TO3-Biotin, and ThT to RNA Mango in the presence of 1 μM Pb^2+^.

## Acknowledgements

We thank Professor Bradley Smith and Hailey Salaberry for providing access to their lab’s fluorimeter. We thank the members of the DasGupta Lab for their valuable feedback on the manuscript. We thank Professor Marya Lieberman and Vikrant Jandev for sharing the salts used for the studies. We thank high school summer students Areej Arif, Jennifer Yang, and Ishita Awasthi for their help during the initial phase of the project. This work was supported by the University of Notre Dame Start Up funds to S.D.

## References

1 J. Yang, X. Li, Z. Xiong, M. Wang and Q. Liu, Environmental pollution effect analysis of lead compounds in China based on life cycle, Int J Environ Res Public Health, DOI:10.3390/ijerph17072184.

2 S. M. C. L. Gioia, M. Babinski, D. J. Weiss, B. Spiro, A. A. F. S. Kerr, T. G. Veríssimo, I. Ruiz and J. C. M. Prates, An isotopic study of atmospheric lead in a megacity after phasing out of leaded gasoline, Atmos Environ, 2017, 149, 70–83.

3 Z. Sun, H. Cao, X. Zhang, X. Lin, W. Zheng, G. Cao, Y. Sun and Y. Zhang, Spent lead-acid battery recycling in China – A review and sustainable analyses on mass flow of lead, Elsevier Ltd, 2017, preprint, DOI: 10.1016/j.wasman.2017.03.007.

4 D. E. Jacobs, R. P. Clickner, J. Y. Zhou, S. M. Viet, D. A. Marker, J. W. Rogers, D. C. Zeldin, P. Broene and W. Friedman, The Prevalence of Lead-Based Paint Hazards in U.S. Housing, 2002, 110.

5 Y. T. Endale, A. Ambelu, G. Sahilu G.B. Mees, and G. Du Laing, Exposure and health risk assessment from consumption of Pb contaminated water in Addis Ababa, Ethiopia, Heliyon, 2021, 7, e07946. 10.1016/j.heliyon.2021.e07946.

6 K. N. Wahyusi, L. I. Utami, S. Aprilio, and N. Fergina, Reduction of Pb and Cr levels in paper industrial Liquid waste with ion exchange method, Journal of Physics: Conference Series, 2020, 1569, 042054 doi: 10.1088/1742-6596/1569/4/042054.

7 S. Zahran, D. Mushinski, S. P. McElmurry and C. Keyes, Water lead exposure risk in Flint, Michigan after switchback in water source: Implications for lead service line replacement policy, Environ Res, 2019, 108928. DOI:10.1016/j.envres.

8 D. Ramírez Ortega, D.F. González Esquivel, T. Blanco Ayala, B. Pineda, S. Gómez Manzo, J. Marcial Quino, P. Carrillo Mora and V. Pérez de la Cruz, Cognitive impairment induced by lead exposure during lifespan: Mechanisms of lead neurotoxicity, MDPI AG, 2021, preprint, DOI: 10.3390/toxics9020023.

9 C. Gundacker, M. Forsthuber, T. Szigeti, R. Kakucs, V. Mustieles, M. F. Fernandez, E. Bengtsen, U. Vogel, K. S. Hougaard and A. T. Saber, Lead (Pb) and neurodevelopment: A review on exposure and biomarkers of effect (BDNF, HDL) and susceptibility, Elsevier GmbH, 2021, preprint, DOI: 10.1016/j.ijheh.2021.113855.

10 L. He, Z. Chen, B. Dai, G. Li and G. Zhu, Low-level lead exposure and cardiovascular disease: the roles of telomere shortening and lipid disturbance, J. Toxicol. Sci. 2018, 43, 11, 623–630.

11 M. L. Adham, Renal Effects of Environmental and Occupational Lead Exposure, Environ Health Perspect, 1997, 105, 928–939. http//ehis.niehs.nih.gov.

12 A. Reuben, M. L. Elliott, W. C. Abraham, J. Broadbent, R. M. Houts, D. Ireland, A. R. Knodt, R. Poulton, S. Ramrakha, A. R. Hariri, A. Caspi and T. E. Moffitt, Association of childhood lead exposure with MRI measurements of structural brain integrity in midlife, JAMA -Journal of the American Medical Association, 2020, 324, 1970–1979.

13 B. P. Lanphear, R. Hornung, J. Khoury, K. Yolton, P. Baghurst, D. C. Bellinger, R. L. Canfield, K. N. Dietrich, R. Bornschein, T. Greene, S. J. Rothenberg, H. L. Needleman, L. Schnaas, G. Wasserman, J. Graziano and R. Roberts, Low-level environmental lead exposure and children’s intellectual function: An international pooled analysis, Environ Health Perspect, 2005, 113, 894–899.

14 P. Levallois, P. Barn, M. Valcke, D. Gauvin and T. Kosatsky, Public Health Consequences of Lead in Drinking Water, Springer, 2018, preprint, DOI: 10.1007/s40572-018-0193-0.

15 S. C. Izah, N. Chakrabarty and A. L. Srivastav, A review on heavy metal concentration in potable water sources in Nigeria: Human health effects and mitigating measures, Springer Netherlands, 2016, preprint, DOI: 10.1007/s12403-016-0195-9.

16 E. Bulska and A. Ruszczyńska, Analytical techniques for trace element determination, Physical Sciences Reviews, DOI:10.1515/psr-2017-8002.

17 Z. ul Nisa, N. A. Ashashi, R. Singhaal, M. Ahmad, R. M. Gomila, A. Frontera and H. N. Sheikh, Selective and efficient detection of Pb2+ in aqueous solution by lanthanoid-organic frameworks bearing pyridine-3,4-dicarboxylic acid and glutaric acid, CrystEngComm, 2023, 25, 2418–2440.

18 M. Sadia, J. Khan, R. Khan, S. W. Ali Shah, A. Zada, M. Zahoor, R. Ullah and E. A. Ali, Trace level detection of Pb2+ ion using organic ligand as fluorescent-on probes in aqueous media, Heliyon, DOI:10.1016/j.heliyon.2024, e41125.

19 S. Suguna, D. Parimala devi, A. Abiram, P. Mukhil sukitha, V. Rajesh kannan, R. Suresh Kumar, A. I. Almansour, K. Perumal, J. Prabhu and R. Nandhakumar, Symmetric and disulfide linked reversible fluorescent organic material: A chemosensor for Pb2+ ion and its applications in real world sample analysis, J Photochem Photobiol A Chem, 2023, 114777. DOI:10.1016/j.jphotochem..

20 C. I. L. Justino, A. C. Freitas, R. Pereira, A. C. Duarte and T. A. P. Rocha Santos, Recent developments in recognition elements for chemical sensors and biosensors, Elsevier B.V., 2015, preprint, DOI: 10.1016/j.trac.2015.03.006.

21 X. Wang, X. Lu, and J. Chen, Development of biosensor technologies for analysis of environmental Contaminants, Trends Environ. Anal. Chem., 2014, 2, 25–32.

22 B. yue Zhang, L. Shi, X. ying Ma, L. Liu, Y. Fu and X. feng Zhang, Advances in the Functional Nucleic Acid Biosensors for Detection of Lead Ions, Taylor and Francis Ltd., 2023, preprint, DOI: 10.1080/10408347.2021.1951648.

23 S. Zhan, Y. Wu, L. Wang, X. Zhan and P. Zhou, A mini-review on functional nucleic acids-based heavy metal ion detection, Elsevier Ltd, 2016, preprint, DOI: 10.1016/j.bios.2016.06.075.

24 J. Li and Y. Lu, A highly sensitive and selective catalytic DNA biosensor for lead ions [9], 2000, preprint, DOI: 10.1021/ja0021316.

25 J. Liu and Y. Lu, A colorimetric lead biosensor using DNAzyme-directed assembly of gold nanoparticles, J Am Chem Soc, 2003, 125, 6642–6643.

26 H. Sun, L. Yu, H. Chen, J. Xiang, X. Zhang, Y. Shia, Q. Yang, A. Guan, Q. Lia, and V. Tang, A colorimetric lead (II) ions sensor based on selective recognition of G-quadruplexes by a clip-like cyanine dye, Talanta, 2015, 136, 210–214. 10.1016/j.talanta.2015.01.027..

27 S. Zhan, Y. Wu, L. Liu, H. Xing, L. He, X. Zhan, Y. Luobc, and P. Zhou, A simple fluorescent assay for lead (II) detection based on lead (II)-stabilized G-quadruplex formation, RSC Advances, 2013, 3, 16962. doi: 10.1039/c3ra42621a.

28 T. Li, E. Wang and S. Dong, Lead(II)-induced allosteric G-quadruplex DNAzyme as a Colorimetric and chemiluminescence sensor for highly sensitive and selective Pb2+ detection, Anal Chem, 2010, 82, 1515–1520.

29 F. Li, L. Yang, M. Chen, Y. Qian and B. Tang, A novel and versatile sensing platform based on HRP-mimicking DNAzyme-catalyzed template-guided deposition of polyaniline, Biosens Bioelectron, 2013, 41, 903–906.

30 S. Dasgupta, S. A. Shelke, N. S. Li and J. A. Piccirilli, Spinach RNA aptamer detects lead(ii) with high selectivity, Chemical Communications, 2015, 51, 9034–9037.

31 J. Mohanty, N. Barooah, V. Dhamodharan, S. Harikrishna, P. I. Pradeepkumar and A. C. Bhasikuttan, Thioflavin T as an efficient inducer and selective fluorescent sensor for the human telomeric G-quadruplex DNA, J Am Chem Soc, 2013, 135, 367–376.

32 T. Bradford, P. A. Summers, A. Majid, P. S. Sherin, J. Y. L. Lam, S. Aggarwal, J. B. Vannier, R. Vilar and M. K. Kuimova, Imaging G-Quadruplex Nucleic Acids in Live Cells Using Thioflavin T and Fluorescence Lifetime Imaging Microscopy, Anal Chem, DOI:10.1021/acs.analchem.4c04207.

33 L. Sjekloća and A.R. Ferré-D’Amaré, Binding between G Quadruplexes at the Homodimer Interface of the Corn RNA Aptamer Strongly Activates Thioflavin T Fluorescence, Cell Chem Biol, 2019, 26, 1159-1168.e4.

34 A. R. De La Faverie, A. Guédin, A. Bedrat, L. A. Yatsunyk and J. L. Mergny, Thioflavin T as a fluorescence light-up probe for G4 formation, Nucleic Acids Res, DOI:10.1093/nar/gku111.

35 G. S. Filonov, J. D. Moon, N. Svensen and S. R. Jaffrey, Broccoli: Rapid selection of an RNA mimic of green fluorescent protein by fluorescence-based selection and directed evolution, J Am Chem Soc, 2014, 136, 16299–16308.

36 E. V. Dolgosheina, S. C. Y. Jeng, S. S. S. Panchapakesan, R. Cojocaru, P. S. K. Chen, P. D. Wilson, N. Hawkins, P. A. Wiggins and P. J. Unrau, RNA Mango aptamer-fluorophore: A bright, high-affinity complex for RNA labeling and tracking, ACS Chem Biol, 2014, 9, 2412– 2420.

37 J. Wu, N. Svensen, W. Song, H. Kim, S. Zhang, X. Li and S. R. Jaffrey, Self-Assembly of Intracellular Multivalent RNA Complexes Using Dimeric Corn and Beetroot Aptamers, J Am Chem Soc, 2022, 144, 5471–5477.

38 K. Y. S. Kong, S. C. Y. Jeng, B. Rayyan and P. J. Unrau, RNA Peach and Mango: orthogonal two-color fluorogenic aptamers distinguish nearly identical ligands, DOI:10.1261/rna.

39 A. Autour, S. C. Y. Jeng, A. D. Cawte, A. Abdolahzadeh, A. Galli, S. S. S. Panchapakesan, D. Rueda, M. Ryckelynck and P. J. Unrau, Fluorogenic RNA Mango aptamers for imaging small non-coding RNAs in mammalian cells, Nat Commun, DOI:10.1038/s41467-018-02993-8.

40 E. V. Dolgosheina, S. C. Y. Jeng, S. S. S. Panchapakesan, R. Cojocaru, P. S. K. Chen, P. D. Wilson, N. Hawkins, P. A. Wiggins and P. J. Unrau, RNA Mango aptamer-fluorophore: A bright, high-affinity complex for RNA labeling and tracking, ACS Chem Biol, 2014, 9, 2412– 2420.

41 R. del Villar-Guerra, J. O. Trent and J. B. Chaires, G-Quadruplex Secondary Structure Obtained from Circular Dichroism Spectroscopy, Angewandte Chemie - International Edition, 2018, 57, 7171–7175.

