## Supplementary Information for "RNA Mango-based Sensors for Lead"

**Table of Contents**


| | | |
|---|---|---|
| Table S1 | Dyes used for the studies, their source, spectral properties, and chemical structures | S3 |
| Table S2 | Oligonucleotide sequences used for the studies | S4 |
| Figure S1 | Crystal structure-derived secondary structures of RNA aptamers screened for $Pb^{2+}$ detection | S5 |
| Figure S2 | Fluorescence spectra of the RNA aptamer-dye pairs in the absence and presence of $Pb^{2+}$ | S6 |
| Figure S3 | The linear region of the fluorescence intensity vs. $Pb^{2+}$ concentration plot used to calculate the limit of detection. | S7 |
| Figure S4 | Fluorescence signal emitted by Mango sensors with TO1-Biotin, TO3-Biotin, and Thioflavin-T (ThT) with increasing concentration of $Pb^{2+}$ | S8 |
| Figure S5 | Dye binding to RNA Mango in the presence of $Pb^{2+}$ | S9 |
| Figure S6 | Fluorescence signal emitted by Mango sensors with TO1-Biotin, TO3-Biotin, and Thioflavin-T (ThT) with different metal ions | S10 |

**Supplementary Tables**

**Table S1**. **Dyes used for the studies, their source, spectral properties, and chemical structures.**

| **Dye/Aptamer** | **$\lambda_{ex}$ (nm)** | **$\lambda_{em}$ (nm)** | **Commercial Source** | **Structure** |
|---|---|---|---|---|
| TO1-3PEG-Biotin/Mango | 510 | 535 | Applied Biological Materials (abm) Inc. | 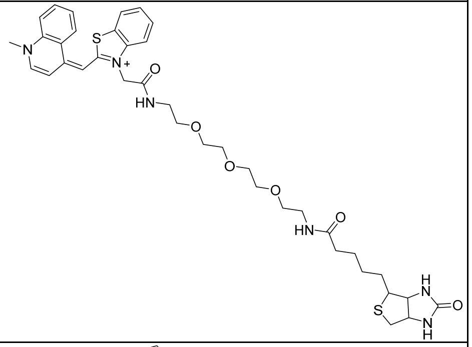 |
| TO3-3PEG-Biotin/Mango | 637 | 658 | Applied Biological Materials (abm) Inc. | 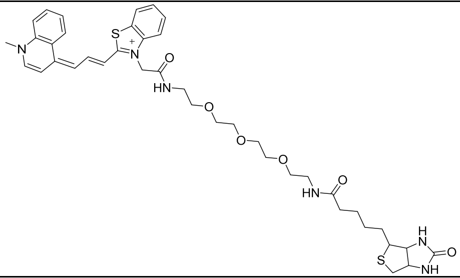 |
| Thioflavin-T (ThT) /Mango | 450 | 484 | Millipore Sigma | 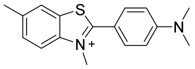 |
| DFHBI-1T/Broccoli | 472 | 507 | Lucerna Technologies | 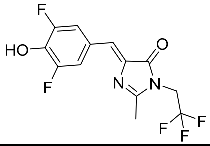 |
| DFHO/Beetroot | 473 | 551 | Lucerna Technologies | 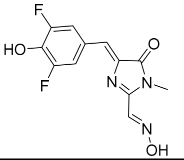 |

**Table S2. Oligonucleotide sequences used in this study.** Nucleotides that constitute the G-quadruplex are depicted in blue. Mutations to G-quadruplex nucleotides are depicted in red.

| Aptamer Name | Oligo sequences |
|---|---|
| Broccoli | 5'-GAGACGGUCGGGUCCAGAUAUUCGUAUCUGUCGAGUAGA GUGUGGGCUC-3' |
| Peach | 5'- GCCCUGAAGGAUGUGGGGGAGGGAGAAGGGC -3' |
| Beetroot | 5'- GGCAGAGGUGGGUGGUGUGGAGGAGUAUCUGU -3' |
| Mango | 5'- GUACGAAGGAAGGUUUGGUAUGUGGUAUAUUCGUAC -3' |
| Mango-G8A (also, MutI) | 5'- GUACGAAAGAAGGUUUGGUAUGUGGUAUAU UCGUAC -3' |
| Mango-G9A (also, MutII) | 5'- GUACGAAGAAAGGUUUGGUAUGUGGUAUAUUCGUAC -3' |
| Mango-G8G9A (also, MutIII) | 5'- GUACGAAAAAAGGUUUGGUAUGUGGUAUAUUCGUAC -3' |

### Supplementary Figures

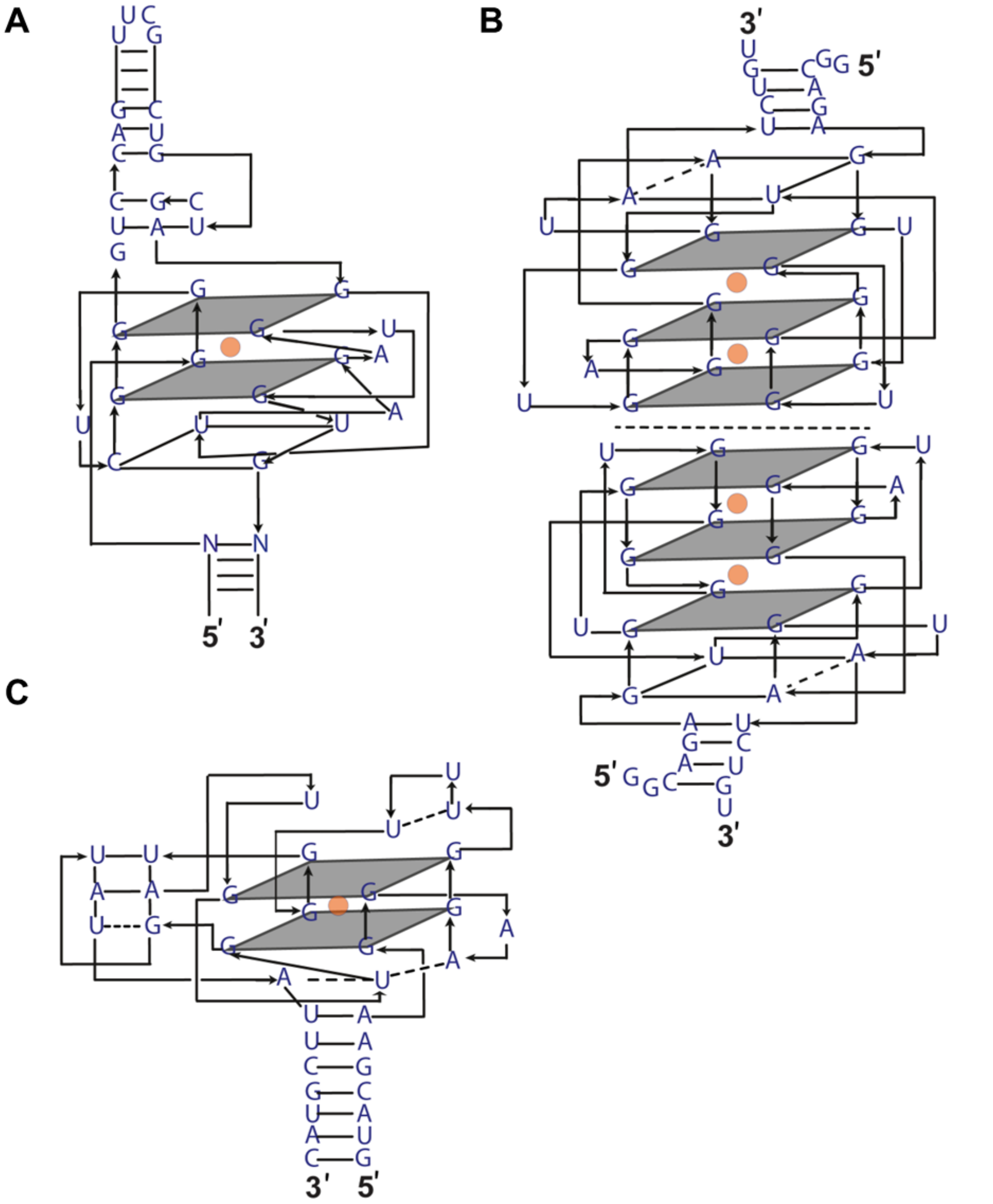


**Figure S1. Crystal structure-derived secondary structures of the RNA aptamers screened for $Pb^{2+}$ detection.** (A) Broccoli, (B) Beetroot, (C) Mango. Broccoli and Mango form a two-layered G-quadruplex, while each monomer of the Beetroot homodimer contains a three-layered G-quadruplex. G-quartets are depicted in grey, and $K^+$ ions are shown as orange circles. The structure of Peach is not shown as no crystal structure is available.

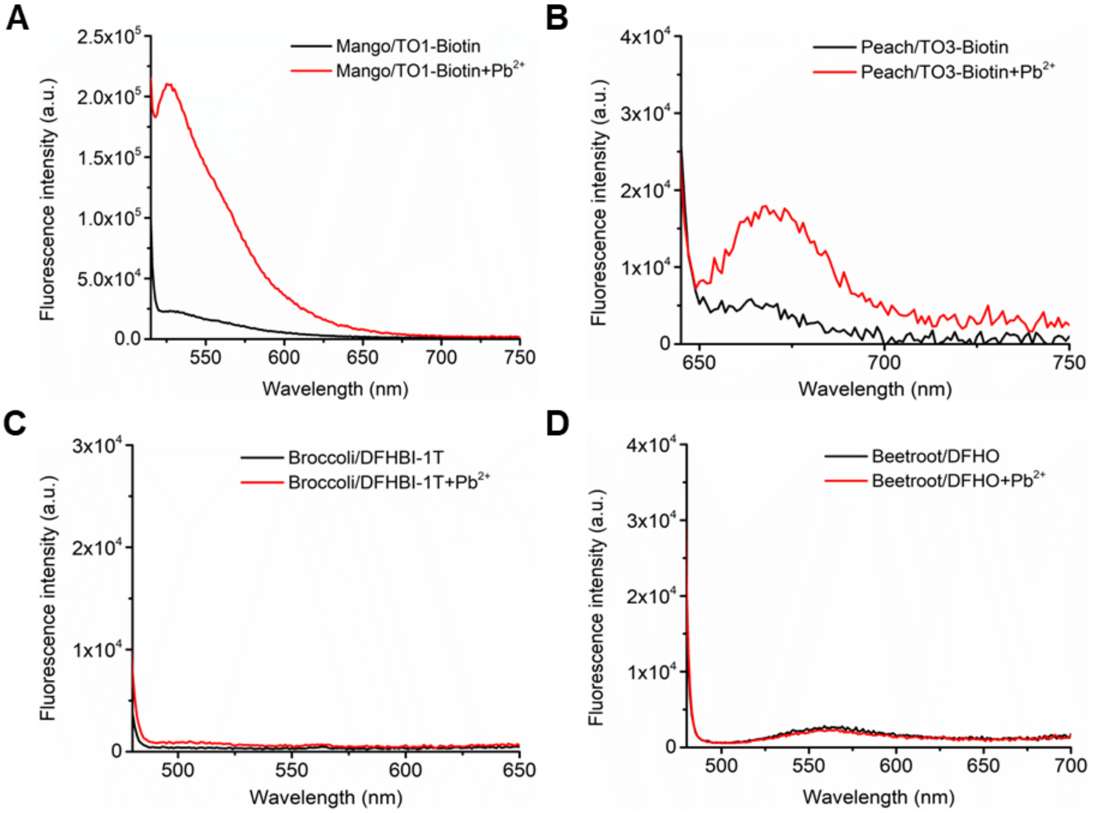


**Figure S2. Fluorescence spectra of the RNA aptamer-dye pairs in the absence and presence of $Pb^{2+}$.** Fluorescence spectra of (A) Mango/TO1-Biotin, (B) Peach/TO3-Biotin, (C) Broccoli/DFHBI-1T, and (D) Beetroot/DFHO in the absence (black) and presence of 1 μM $Pb^{2+}$ (red). Mango/TO1-Biotin and Peach/TO3-biotin showed fluorescence enhancements with $Pb^{2+}$. Experiments were performed at pH 8 and 5 mM $Mg^{2+}$. Concentrations of RNA and dye were 300 nM and 3 μM, respectively.

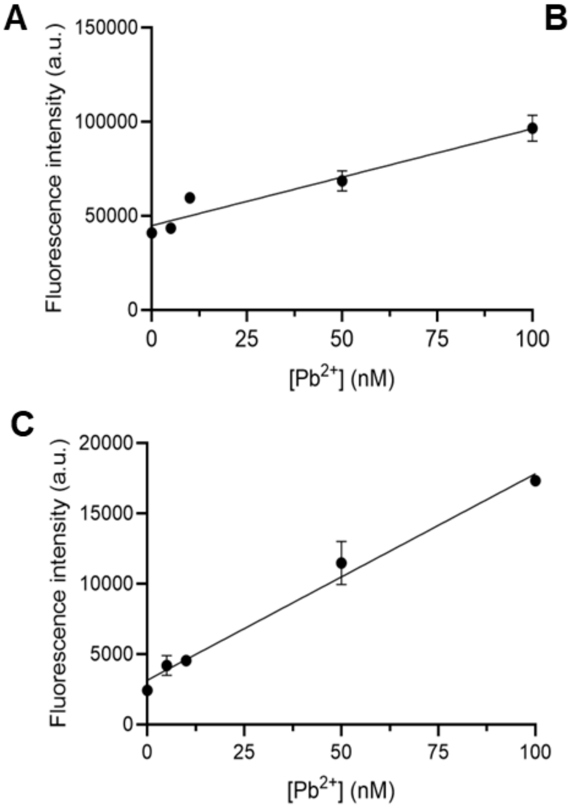


**Figure S3. The linear region of the fluorescence intensity vs. $Pb^{2+}$ concentration plot used to calculate the limit of detection.** Mango fluorescence *versus* $Pb^{2+}$ concentration for (A) TO1-Biotin ($R^2$ = 0.91), (B) TO3-Biotin ($R^2$ = 0.91), and (C) ThT ($R^2$ = 0.97). Signal increases linearly with $Pb^{2+}$ concentration in the range of 5 nM-100 nM. Experiments were performed at pH 8 and 5 mM $Mg^{2+}$. Concentrations of RNA and dye were 100 nM and 1 µM, respectively. See Figure 4A-C for the full plot. Error bars correspond to the standard deviation of three independent experiments.

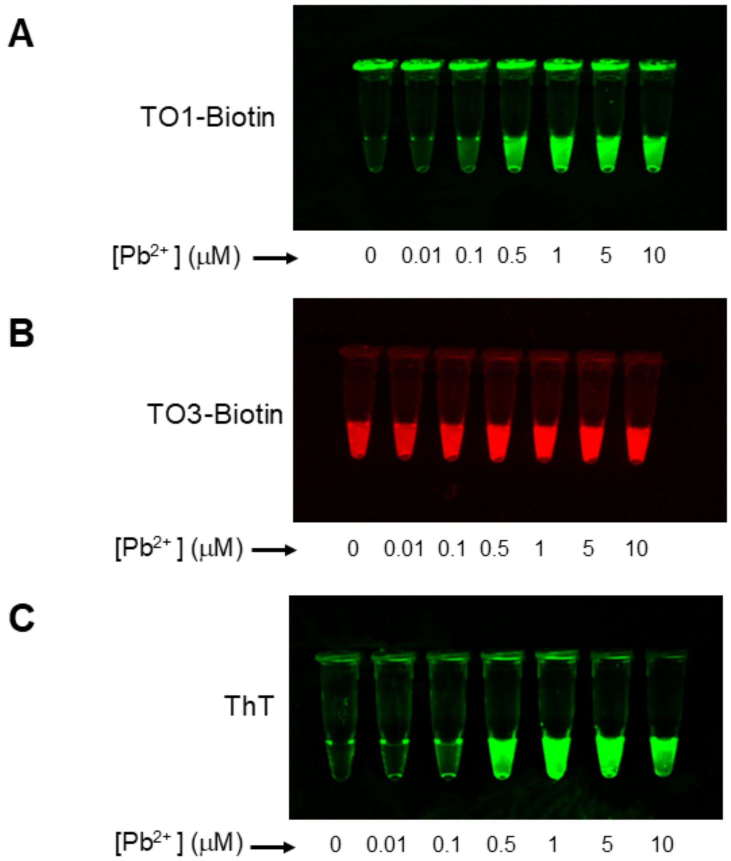


**Figure S4. Fluorescence signal emitted by Mango sensors with TO1-Biotin, TO3-Biotin, or Thioflavin-T (ThT) dyes with increasing concentration of $Pb^{2+}$.** Concentration-dependent fluorescence enhancement is observed for TO1-Biotin and Thioflavin-T. Experiments were performed at pH 8 and 5 mM $Mg^{2+}$. Concentrations of RNA and dye were 100 nM and 1 μM, respectively. 10 nM – 10 μM $Pb^{2+}$ were used.

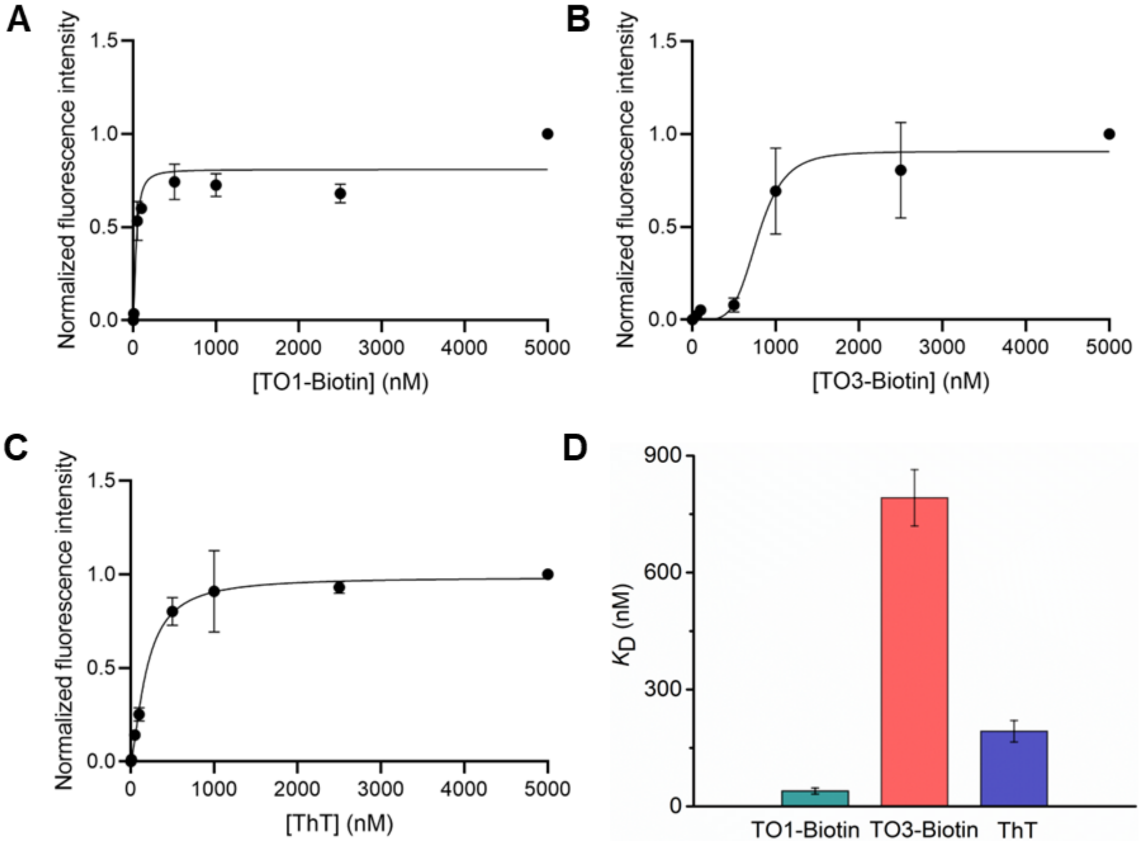


**Fig. S5. Dye binding to RNA Mango in the presence of $Pb^{2+}$.** Fluorescence signal exhibited by RNA Mango in response to increasing concentrations of (A) TO1-Biotin, (B) TO3-Biotin, and (C) ThT in the presence of $Pb^{2+}$. Fluorescence increases with increasing dye concentrations from 5 nM – 5 μM. (D) RNA Mango binds to TO1-Biotin and ThT with significantly higher affinities than to TO3-Biotin. Experiments were performed at pH 8 and 5 mM $Mg^{2+}$. Concentrations of RNA and $Pb^{2+}$ were 100 nM and 1 μM, respectively. Error bars correspond to the standard deviation of three independent experiments.

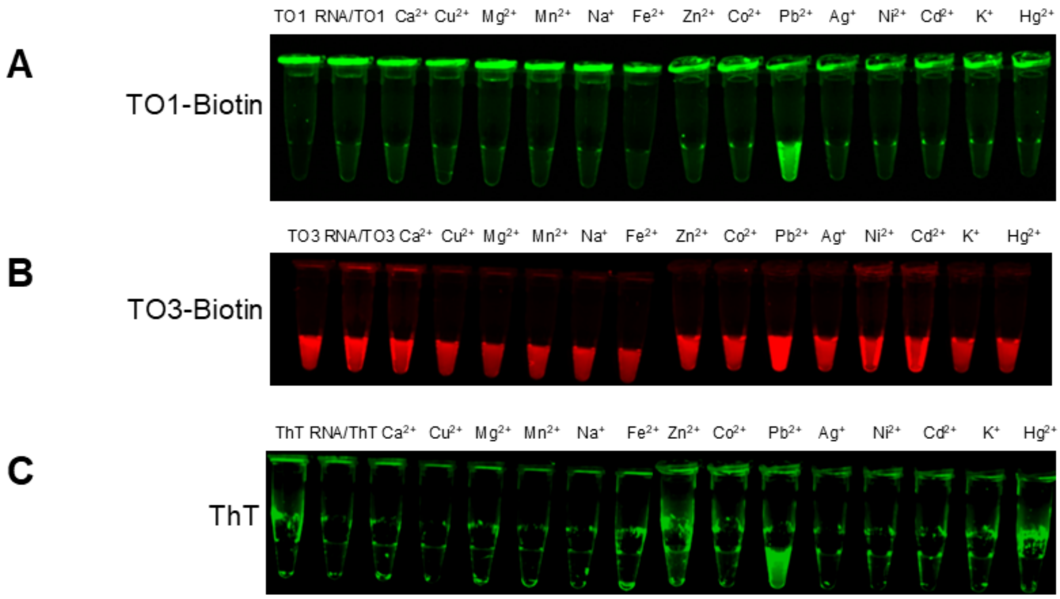


**Figure S6**. **Fluorescence signal emitted by Mango sensors with TO1-Biotin, TO3-Biotin, and Thioflavin-T (ThT) with different metal ions**. A selective fluorescence signal is observed only in the presence of $Pb^{2+}$ ions. Experiments were performed at pH 8 and 5 mM $Mg^{2+}$. Concentrations of RNA and dye were 100 nM and 1 μM, respectively in all samples containing metal ions, which were added to 1 μM.
